# The interaction between aging and DNA damage influences the accumulation of APP in Alzheimer’s disease

**DOI:** 10.1101/2025.09.07.674780

**Authors:** Eliana Brenner, Gnanesh Gutta, Fiona Stauffer, Jay Mehta, Rody Kingston, Michael Bagnell, Starr Welty, Natalie Dixon, Elizabeth Zinzell, Karl Herrup, Fulin Ma

## Abstract

DNA damage is a major driver of the aging process, and the progression of Alzheimer’s disease (AD) worsens the situation. Neurons of the AD brain show significantly elevated levels of DNA damage compared to controls. AD is also associated with altered processing of the amyloid precursor protein (APP), best known for its Aβ proteolytic peptide fragment. In the current work we find that these three elements – age, DNA damage, and APP – are associated with each other. We present evidence from the mouse brain supporting the hypothesis that DNA damage drives increased levels of APP. Using the TUNEL reaction to track DNA damage, we show that TUNEL staining increases significantly with age (6 months to 24 months, 8 females and 14 males total in this study), in lockstep with intracellular APP. To separate correlation from causality we analyzed tissue from mice genetically deficient in ATM (ataxia-telangiectasia mutated), a protein kinase that promotes DNA damage repair. Compared to 6-month wild type mice, the increased TUNEL signal in neurons of 6-month *Atm^−/−^* animals was equivalent to that seen in 24-month wild type. Significantly, the APP signal in the *Atm^−/−^* cells was also increased, and this correlation was found in both neuronal and non-neuronal cells. These results suggest that the loss of genomic integrity associated with aging is a cellular stressor that increases the levels of APP and thus increases vulnerability to the pathogenesis of AD.

**SIGNIFICANCE STATEMENT:** Our work links the genetics of Alzheimer’s disease (dominant disease-causing mutations in the APP gene) with DNA damage and aging – two factors known as risk factors in AD. The evidence suggests that the enhanced DNA damage associated with the normal aging process, increases the levels of APP thus potentially contributing to Aβ production and plaque formation.

## INTRODUCTION

Age is the single greatest risk factor for several different neurodegenerative diseases, especially dementing illnesses such as Alzheimer’s disease (AD). It follows, therefore, that understanding the biology of brain aging should be part and parcel of any therapeutic approach to slowing or halting the progress of age-related dementias. Unfortunately, aging remains a mysterious process whose biological features remain incompletely understood. One important clue that has guided research over the past decades is that the process of aging is inextricably linked with the loss of genomic integrity. Despite the robust nature of the correlation, however, the mechanisms tying aging to DNA damage are complex and have yet to be fully discovered. DNA damage accumulates over the lifespan (Lu et al., 2004), and several lines of evidence suggest that this damage is itself one of the primary causes of organismal aging (McKinnon, 2013; Chow and Herrup, 2015; Tse and Herrup, 2017; Yousefzadeh et al., 2021). For individual cells, the data are even more compelling. Cellular senescence is a state of cellular physiology that is nearly synonymous with aging. The number of senescent cells increases with age in virtually every tissue (Chinta et al., 2015), and once a cell senesces, it permanently loses its capacity for cell division and enters a state known as the senescence-associated secretory phenotype. There are several conditions that cause cells to senesce, but one of the most notable is the induction of DNA damage (Campisi, 2013; van Deursen, 2014; Schwab et al., 2019; Kim et al., 2020). Thus, the loss of genomic integrity likely drives both organismal and cellular aging.

The question then becomes how does age and/or DNA damage affect the specific symptoms of clinically distinct neurological conditions? For example, both Parkinson’s and Alzheimer’s are age-related dementias, and both have increased DNA damage as one of their characteristics (Coppede and Migliore, 2015), yet their clinical symptoms and neuropathology are quite different. There is ample evidence that DNA damage increases in AD compared to age-matched controls (Suberbielle et al., 2013; Suberbielle et al., 2015; Lin et al., 2020; Gutta et al., 2024; Sharma et al., 2025), and that the damage appears early in the progression of the disease (Shanbhag et al., 2019). To explain how the response to this damage differs from that found in a disease such as Parkinson’s, one might speculate that there is a disease-specific pattern to the accumulation of new DNA damage. This seems highly unlikely since, while it may not be totally random, new DNA damage caused either by endogenous cellular insults, such as free radicals, or by exogenous factors such as environmental pollutants should not be specific to any one gene sequence. Thus, it becomes important to know whether the DNA damage associated with AD is different from that found in Parkinsons. The answer may be hidden in the fact that while DNA damage may not be specific to a particular gene sequence, DNA damage repair could be selective. This leads to the speculation that it is the response to DNA damage that is the key factor differentiating age-related dementias.

The proteins known as α-synuclein and amyloid precursor protein (APP) are associated with Parkinson’s and Alzheimer’s, respectively. Both have proteolytic fragments that aggregate into unusual inclusions that are seen as hallmarks of the two diseases. Less appreciated is that both proteins play important roles in the process of DNA repair (Schaser et al., 2019; Revol et al., 2023). In the case of APP, it is not the full-length transmembrane protein, but rather the APP intracellular domain (AICD) that is the active agent. The AICD associates with Fe65 and migrates to the nucleus where it joins with TRRAP and the histone acetylase, TIP60, to induce chromatin modifications that facilitate DNA repair (Stante et al., 2009; Revol et al., 2023). This pathway is integrated into the DNA damage response since inducing DNA damage increases the γ-secretase activity that is needed to generate the AICD (Jin et al., 2007; Jin et al., 2008), although exactly how remains unclear.

Our laboratory has been focused on the role of the full-length APP protein and its sensitivity to neuronal stresses. In vivo, axonal damage such as that incurred during traumatic brain injury can rapidly increase the expression of the *APP* gene in sheep brain (Van den Heuvel et al., 1999a). In vitro, even a mild excitatory stimulus (1 µM glutamate) is sufficient to increase neuronal *App* mRNA and APP protein by several fold (Ma et al., 2023). This led us to ask whether DNA damage might serve as another neuronal stress that would induce *App* expression. We opted to approach this question *in vivo* by inhibiting DNA damage repair rather than inducing exogenous DNA damage. We took advantage of mice that carried engineered mutations in the *Atm* gene (Xu et al., 1996). ATM (ataxia-telangiectasia mutated) is a member of the PI3-kinase family that is involved in many cellular activities including the non-homologous end-joining process of DNA repair. In the absence of ATM activity, the rate and efficiency of DNA repair is reduced (Lee and McKinnon, 2007; Goodarzi et al., 2008). In addition, the increased APP from DNA damage accelerates APP processing and contributes to Aβ generation (Almenar-Queralt et al., 2014; Welty et al., 2022). As a second approach, we used the increase in DNA damage that is found during the natural aging process. We report here that both means of increasing DNA damage significantly increase cellular APP levels. The findings thus offer important new insights into the linkages among aging, DNA damage, and dementing illnesses such as Alzheimer’s disease.

## MATERIALS AND METHODS

### Mice

Wild type (WT) C57BL/6J mice and breeding stock of *Atm^−/−^* (B6;129S4-*Atm^tm1Bal^*/J) were purchased from The Jackson Laboratory then bred and aged in our colony. The *Atm* stock was maintained by backcrossing heterozygous (+/*Atm^-^*) animals to wild type C57BL/6J mates. This breeding scheme is required as *Atm* homozygous animals are sterile. Homozygous (*Atm^−/−^*) animals were produced by intercrossing heterozygotes. The genotype of all *Atm* mice were confirmed by PCR using the protocol listed on The Jackson Laboratory website. Each animal was genotyped twice to confirm their genetic assignment. A total of 8 females and 14 males were used in this study.

### Histology

Mice were injected with a mixture of 100 mg/kg ketamine hydrochloride (Covetrus, 118374) and 20 mg/kg xylazine (Covetrus 1XYL006). The mice were then perfused with 20 mL 0.9% saline at 2.5 mL/min. After perfusion, the brains were dissected free of the skull. The left hemispheres were dissected into cortex and cerebellum and stored immediately into −80 °C freezer for protein analysis. The right hemispheres were immersion-fixed in 4% PFA at 4 °C for 24 hours. After the fixation, the brains were transferred to 30% sucrose and stored at 4°C for 24 hours. The following day the brains were bisected on the midline, embedded in optimal cutting temperature (OCT) gel (Fisher Scientific, 23-730-571) and stored at −80 °C until use. Sagittal cryostat sections were taken at 10 μm and mounted on SuperFrostPlus glass slides (Fisher Scientific, 12-550-15).

### TUNEL

DNA strand breaks were detected and measured with a Click-IT TUNEL (terminal deoxynucleotidyl transferase dUTP nick end labeling) Alexa Fluor 647 imaging assay (Thermo Fisher Scientific, C10247) following the manufacturer’s instructions. Briefly, the tissue samples were washed with phosphate buffered saline (PBS) three times followed by a permeabilization with 0.3% Triton X-100 in PBS for 20 minutes. The tissue slides were next incubated with 200µL of the terminal deoxynucleotidyl transferase (TdT) reaction buffer for 10 minutes under room temperature. Tissues were then incubated in 200 µL of TdT cocktail for 1 hour at 37 °C in a humidity incubator. Cells were then washed twice in 3% bovine serum albumin (BSA) in 1 X PBS for 5 minutes, then incubated with Click-IT reaction buffer with additive for 30 minutes in the humidity incubator. The tissue slides were then washed 3 times in PBS before proceeding to antibody staining where the slides were protected from light for the rest of the experiment.

### Immunohistochemistry

Antibody staining was performed after the TUNEL reaction was complete. Tissue sections were rinsed in the dark with PBS three times for 5 minutes each. Sections were outlined with an oil pen to prevent dispersion of the reagents and define the region of tissue to be stained. Sections were then incubated for one hour at room temperature with 0.3% Triton X-100 in PBS (PBST) with 5% donkey serum to block nonspecific binding. After blocking, the sections were incubated overnight with primary antibodies in 5% donkey serum in PBST. All incubations were carried out at 4 °C in a humidified chamber. The primary antibodies and their working dilutions are listed in Table I. The following day, the mouse tissue sections were rinsed with PBS three times for five minutes each, then incubated for one hour at room temperature with secondary antibodies diluted 1:500 in 5% donkey serum in PBST. The sections were rinsed three times with PBS for five minutes each at room temperature then counterstained with DAPI diluted to 1:100 in PBST for 10 minutes followed by two final five-minute PBS rinses. The sections were mounted under coverslips using Hydromount.

**TABLE I.** Primary antibodies.

| **Primary antibodies** |  |  |  |  |  |
| --- | --- | --- | --- | --- | --- |
| *Antibodies* | *Manufacturer* | *Immunogen* | *Catalog #* | *RRID#* | *Dilution* |
| γ-H2AX | Abcam | Proprietary | ab26350 | AB_470861 | 1:250 |
| NeuN-Cy3 | Millipore | Fox-3 | ABN78C3 | AB_11204707 | 1:250 |
| APP-N-KO | Abcam | Proprietary | ab126732 | AB_11131727 | 1:500 |
| APP-C | Abcam | NPXY motif of human APP | ab32136 | 960 AB_2286 | 1:400 (IF)<br>1:1000 (WB) |
| Neurofilament (smi-312) | Biologend | NF-M+H subunits | 837904 | AB_2566782 | 1:1000 |
| 6E10 | Sigma | β-amyloid peptide | 39320 | AB_662798 | 1:800 (IF) |
| GAPDH | Sigma-Aldrich | N-terminal cysteine residue | G9545 | AB_796208 | 1:1000 (WB)<br>1:1000 |

### Western Blots

Lysates were homogenized in ice-cold RIPA buffer (EMD Millipore, R0278-50ML) with 1× PhosSTOP phosphatase inhibitor mixture (Roche Applied Science, 4906845001) and 1 × complete protease inhibitor cocktail (Thermo Fisher Scientific, 78429). The homogenate was then centrifuged at 4 °C for 20 minutes at 20,000 rcf. The supernatant containing protein was collected. Total protein concentrations of the lysates were determined by Bicinchoninic Acid (BCA) assay (Thermo Fisher Scientific, 23227). Samples were prepared for Western Blot using a lysate dilution that would yield 25-50 µg total protein with 4% β-mercaptoethanol in 2 × Novex Tris-Glycine SDS Sample Buffer (Life Technologies, LC2676). Samples were heated to 90°C for 10 minutes., then run on 4%-20% Novex Wedge Well Tris-Glycine mini gels (Thermo Fisher Scientific, XP04205BOX) in 1 × Tris/Glycine/SDS running buffer (Thermo Fisher Scientific, BP13414) at 80 V for 10 minutes, then 100 V for 1 hour. The protein was transferred to a nitrocellulose membrane using the Invitrogen Power Blotter System (Thermo Fisher Scientific, PB0012) and 80% pre-made transfer buffer stock (5.85 g glycine, 11.6 g Tris, 0.375 g SDS, milliQ water to 1 L) with 20% ethanol at a constant 25 V for 45 minutes. [For MAP2 and 53BP1, protein was transferred to PVDF membrane activated in methanol using 80% pre-made transfer buffer stock (144 g glycine, 30.3 g Tris base, milliQ water to 1 L) with 20% methanol at a constant 80 mA for 24 hours in 4 °C on ice.] Blots were stained with Ponceau (5% acetic acid, 0.1% Ponceau stain) for five minutes to verify total protein transfer. Blots were then blocked in TBST with 5% milk for one hour and immunoblotted overnight at 4°C using the primary antibodies from Table I in 5% milk and TBST. Blots were washed three times with TBST for ten minutes. each, then probed with appropriate secondary antibodies (anti-Mouse IgG Horseradish Peroxidase HRP or anti-Rabbit IgG HRP) in a 1:10000 dilution in 5% milk and TBST for 1 hour at room temperature. Following secondary antibody probing, blots were washed three times for ten minutes. in TBST, then washed once for ten minutes. in TBS. Blots were imaged using ECL substrate (Perkin Elmer, NEL 105001) or SuperSignal West Femto Maximum Sensitivity Substrate (Thermo Fisher Scientific, 34096) in the Bio-Rad ChemiDoc.

### Image analysis

The slides were viewed on either a fluorescence microscope (Nikon, eclipse Ni.) or a confocal microscope (Nikon, AR1). Images of the mouse tissue sections were captured from the deeper layers of frontal cortex of the mouse. Two images were taken per section. We used QuPath (version 0.6.0), an open-source software for digital pathology, for the quantification of the mouse tissue section images. We adjusted the parameters of the positive cell detection software to detect DAPI positive nuclei. We then utilized a single measurement classifier gating on nuclear size supplemented with manual deletions to select neurons based on their DAPI staining and nuclear area. This was followed by utilizing the extension function of QuPath to expand the DAPI-defined nucleus by 4 μm in all directions from its center of gravity. After subtracting the nucleus, we define the remaining area as the “pseudoplasm”. The pseudoplasm partially captures the perinuclear cytoplasm, but due to the non-circular shape of a neuron the software makes both Type I and Type II errors when capturing the perinuclear cytoplasm. Specifically portions of the pseudoplasm will not contain true neuronal cytoplasm (Type I) and virtually all of the neurites extending beyond the 4 µm extension were excluded even when well stained (Type II). The data set was exported as a .csv file and sorted with Excel (version 16.92 [24120731]).

Using FIJI (software version 2.14.0/1.54g), five background regions of interest were randomly selected across any given image where no cells were detected. This was done for the TUNEL and APP channels for every image obtained from the confocal microscope. The intensity of these randomly selected background regions was averaged and used as a background measurement. The TUNEL and APP channel intensities for detected cells were then divided by the respective background measurements to obtain a ratio.

Protein bands from Western Blot imaged using Bio-Rad ChemiDoc gel imager were quantified with FIJI (software version 2.14.0/1.54g). The area selected for quantification included entire individual bands. Intensity density was measured followed by normalization with the respective loading controls.

### Statistical analysis

Data were expressed as mean ± SEM for each group. Statistical analysis was performed using GraphPad Prism software [version 10.4.1 (532)]. The statistical significance of changes in different groups was evaluated by unpaired Student’s t test and one-way ANOVA using GraphPad Prism software. P values <0.05 were considered to be significant.

## RESULTS

### DNA damage increases with age

To assess the levels of DNA damage on a cell-by-cell basis, we considered several alternative methods: immunostaining for DNA repair proteins, such as γ-H2AX or 53BP1, or histochemical staining using the TUNEL reaction (Terminal deoxynucleotidyl transferase dUTP nick end labeling) which labels free 3’ DNA ends such as those found in double strand breaks or single strand nicks (Gavrieli et al., 1992). We chose TUNEL as our metric of DNA damage. We made this choice in part because we used ATM (ataxia-telangiectasia mutated) deficiency as one means of enhancing DNA damage, and the phosphorylation that defines γ-H2AX is normally catalyzed by the ATM kinase (Burma et al., 2001), although the DNA-dependent protein kinase (DNA-PK) can also phosphorylate H2AX (Park et al., 2003; Stiff et al., 2004). Our reservations about the use of γ-H2AX as our DNA damage marker was further supported by double immunostaining. Six-month-old adult wild type mouse brain samples were prepared as described in the Methods section, followed by immunolabeling for Y-H2AX and Click-IT labeling of the polyU tail added during TUNEL (Fig. 1A). We found that the patterns of γ-H2AX and TUNEL staining were similar but nonetheless distinct. There were cells with double-labeling of Y-H2AX and TUNEL (yellow arrows), but also cells with only γ-H2AX (green arrow) or only TUNEL labeling (red arrow). The differences were most likely because Y-H2AX reveals the “history” of DNA damage, while TUNEL reveals current DNA breaks. Y-H2AX staining labels modified H2AX histone proteins that are generated in domains as large as a megabase surrounding the DNA lesion (Savic et al., 2009) and take as long as 24 hours to clear (Suberbielle et al., 2013). The TUNEL reaction, by contrast, reveals only current DNA breaks. Thus, despite our previous work (Gutta et al., 2024) in human material, we chose TUNEL as our primary metric of DNA damage for the current study.

**Figure 1.**
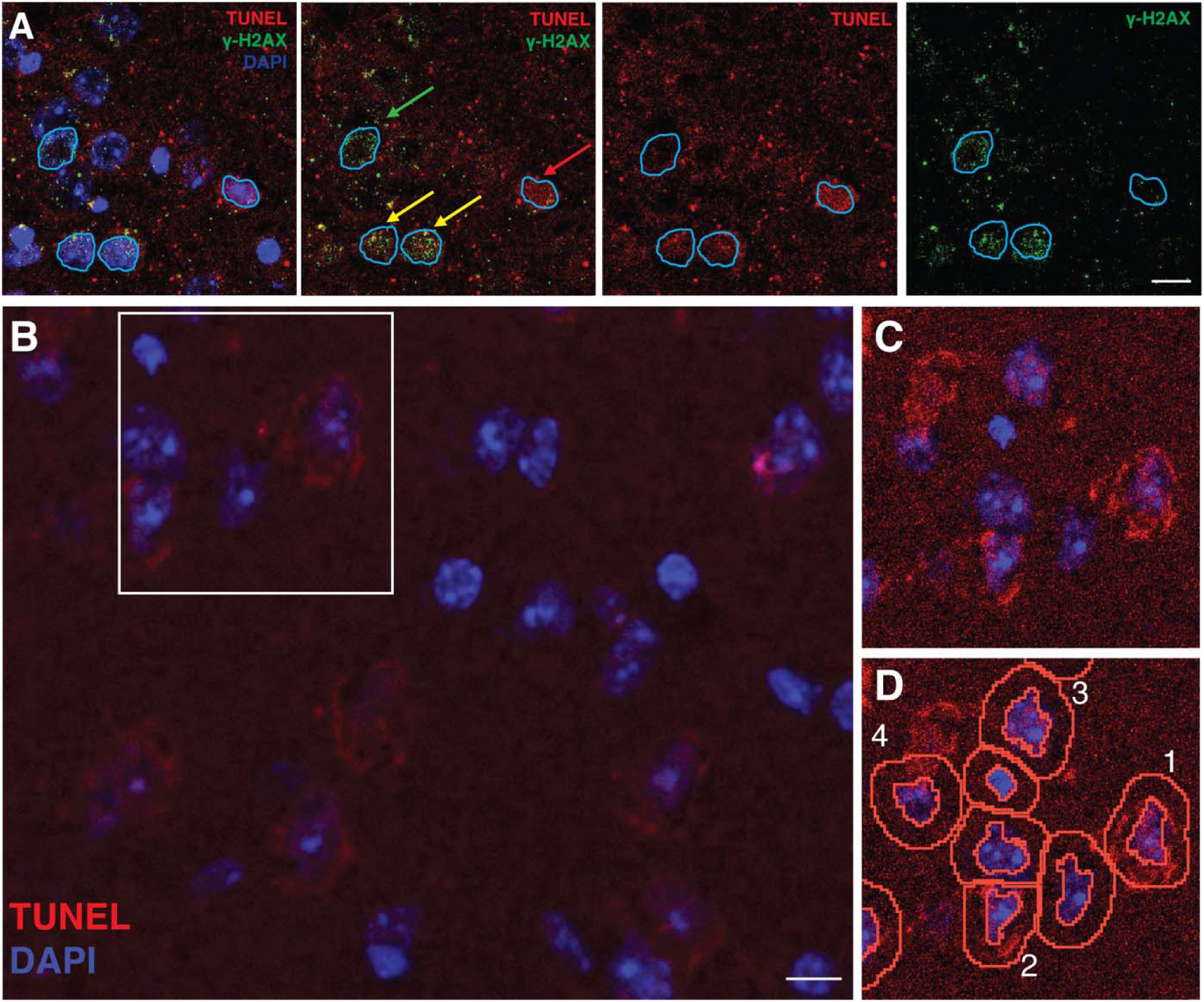
TUNEL detects DNA damage in both the nucleus and cytoplasm of neurons. (A) A 6-month-old mouse cortex stained with Y-H2AX antibody (green), TUNEL (red) and DAPI (blue). Yellow arrow: TUNEL and Y-H2AX double positive. Green arrow: Y-H2AX only. Red arrow: TUNEL only. Scale bar = 10 µm. (B) Neurons of layer V-VI from a 6-month-old mouse prefrontal cortex stained for TUNEL and counterstained for DAPI. Scale bar = 10 µm. (C) High magnification of a small number of TUNEL stained neurons with signal enhanced for clarity. (D) The same image as in (C) with the QuPath detections overlaid. The inner circle is the identified nucleus (based on DAPI only). The outer circle is the “pseudoplasm” defined by extending the nuclear boundary 4 µm from the nuclear center of gravity.

TUNEL staining of the neocortex of six-month old mice yielded images such as those seen in Figure 1B. Six-month-old mice were chosen to represent ‘young’ animals as they are roughly the equivalent of a human in their 20’s. They have a stable neurophysiology and yet have little evidence of age-related DNA damage (Hamilton et al., 2001; Yu et al., 2018). In these young specimens, we found a faint but reliable TUNEL signal in most neurons. While DNA damage would be expected only in the nucleus, note that the TUNEL reaction product is also seen in the cytoplasm, most likely a reflection of cytoplasmic DNA (cytoDNA). As described previously, nuclear DNA damage results in small fragments of genomic DNA being exported to the cytoplasm (Van den Heuvel et al., 1999b; Quek et al., 2017; Song et al., 2019; Song et al., 2021). The images in the figure clearly demonstrate that similar cytoplasmic DNA species (cytoDNA) are found in cortical neurons. QuPath image analysis software (Bankhead et al., 2017) was used to quantify these images. We used DAPI to define the cell nucleus then used the “extension” function of QuPath set to 4 µm to define the “pseudoplasm” (Fig. 1C, D). This particular field was chosen to illustrate both the strengths and weaknesses of this or any image analysis platform. The cells labeled 1, 2, and 3 in Figure 1D are representative of >95% of the cells we measured. The DAPI stained nucleus is well captured by the software (inner circle) and the 4 µm extension captures the cytoplasm of the cells. Cell 4 in Figure 1D illustrates a limitation of the system. The DAPI stained nucleus is well defined, but a nearby cell whose nucleus is not present in the section was defined as the pseudoplasm of cell 4. Because of these limitations, we chose to limit our analysis to DNA damage in the nucleus and forego quantification of the cytoDNA TUNEL signal.

To determine the effects of aging, we next asked whether the neurons in two-year-old mice showed an enhanced TUNEL signal compared with those of younger samples. We first noted that the qualitative appearance of TUNEL labeling was different in the 6-month and 24-month material. In the younger mice, the level of staining was low. This included both cells with larger as well as smaller nuclear size. In the older mice clear TUNEL staining was found in both the nucleus and cytoplasm (Fig. 2A), and the intensity of the TUNEL signal was noticeably higher than that in the 6-month animals (Fig. 1). Using QuPath to quantify the nuclear TUNEL signal as defined by DAPI staining (yellow outlines, Fig. 2Ai and 2Aii), we found that older neurons showed a higher level of TUNEL staining than their younger counterparts (Fig. 2B). Note too that the variance of the sample increased markedly.

**Figure 2.**
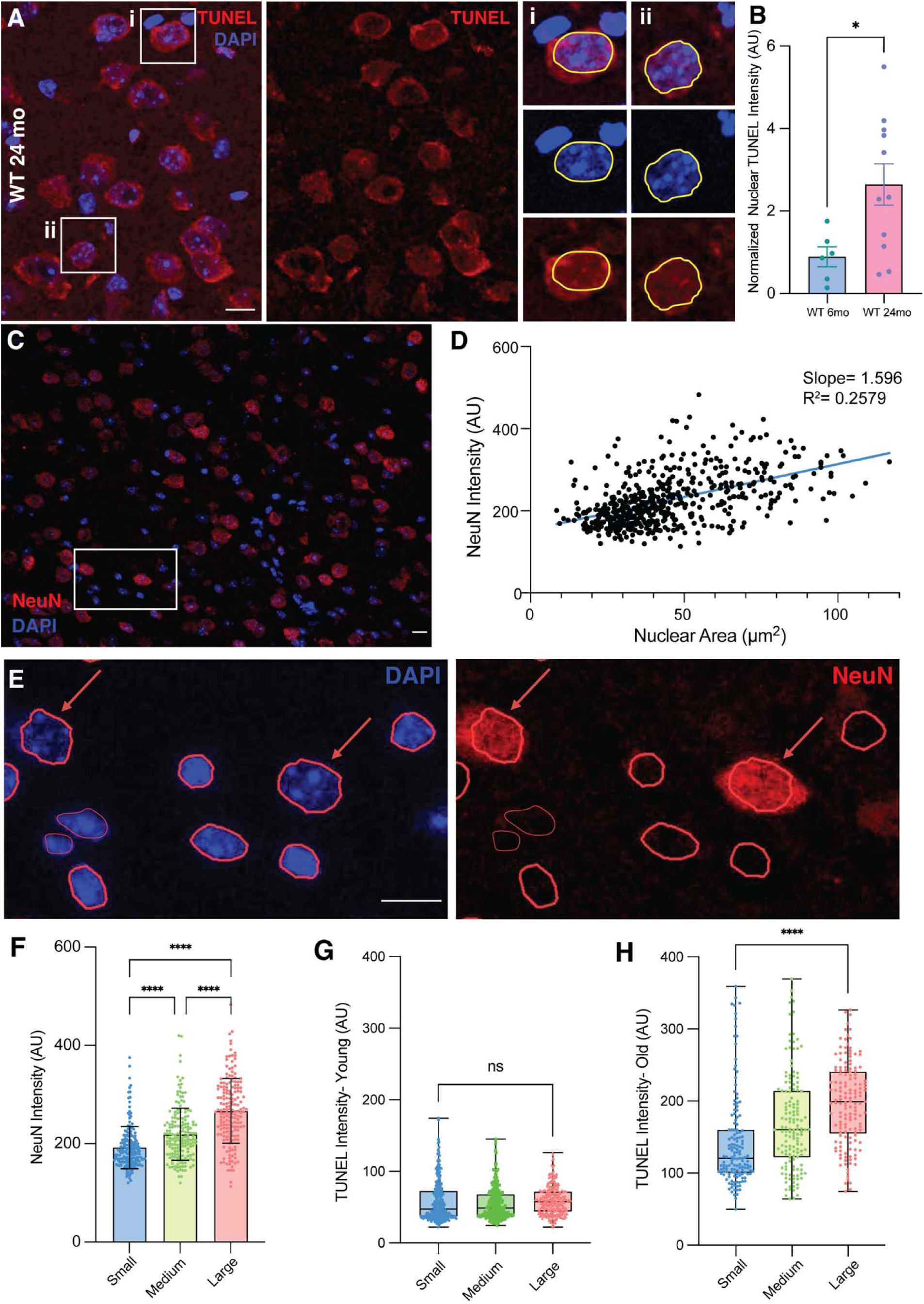
DNA damage accumulates with age in neurons. (A) Neurons of layer V-VI from a 24-month-old mouse prefrontal cortex stained for TUNEL (red) and counterstained for DAPI (blue). Ai and Aii are two representative cells shown enlarged for additional detail. Scale bar = 10 µm. (B) Nuclear TUNEL reaction intensity in 6-month (blue) and 24-month (red) old wild type mice neurons. Each point represents the average of all cells measured in one animal. Error bars represent standard deviation. *, p = 0.0257 by two tailed t-test (t = 14.93, df = 916). (C) Immunolabeling of NeuN (red) and counterstained with DAPI (blue) in a 24-month wild type mouse cortex. Scale bar = 10 µm. (D) A scatter plot showing correlation of nuclear NeuN intensity as a function of the area of the nucleus in (C). (E) Immunolabeling of NeuN (red) of the cortex from the animal shown in the white box from (D). Red circles indicate cell nuclei and red arrows show cells with large nuclei. Scale bar = 10 µm. (F) Plot of NeuN intensity in different cell types. Cells were separated into tertile based on nuclear size. Small: predominantly non-neuronal nuclei; Medium: mixture of cell types; Large: exclusively neuronal nuclei. (One-way ANOVA with multiple comparison. p < 0.0001, df = 350). (G, H) Box and whisker plot of TUNEL intensity in three nuclear size tertiles. (G) In young animals the upper (red) and lower (blue) tertiles were not significantly different by two tailed t-test (p = 0.5764; t = 0.0590, df = 548). (H) In old animals the difference between the upper and lower tertiles was highly significant by two tailed t-test (****, p < 0.0001; t = 7.683, df = 293).

To determine the cell types involved, we correlated nuclear size, shown by DAPI, validated with the intensity of NeuN labeling. NeuN is a nuclear localized neuron-specific marker that is commonly used distinguish neurons from glia (Herculano-Houzel et al., 2006; Mota and Herculano-Houzel, 2014; Erö et al., 2018; Das and Ramanan, 2023). We split the entire population of DAPI defined nuclei into three tertiles based on their area. The tertile with the largest size contained exclusively NeuN- and MAP2-positive cells that we identified as neurons. In the bottom tertile > 98% of the cells were those we identified as non-neuronal (NeuN- and MAP2-negative). As expected, the middle tertile contained a variable mixture of the two. Not surprisingly, cells with larger nuclear area also had stronger NeuN fluorescent labeling (Fig. 2C-F – data from 6-month wild type animals). In this way we determined that in younger mice, the level of DNA damage was low and not significantly different between neurons and non-neurons (Fig. 2G). In the older mice, while all cells had significantly brighter TUNEL staining than cells from younger brains, there was a clear distinction between neurons and non-neurons (Fig. 2H – bottom vs. top tertile). This suggests that DNA damage accumulates to a greater extent in neurons than in non-neuronal cells in old mice.

### APP levels increase with age

Our long-term interests lie in the relationships of aging and DNA damage to the pathogenesis of Alzheimer’s disease. We therefore examined the effects of age on the levels of the amyloid precursor protein (APP) on a cell-by-cell basis. We performed APP immunostaining after completion of the TUNEL reaction and compared the patterns observed. We used antibodies directed against both the C- and N-terminal portions of the full-length protein (APP-C and APP-N, respectively). Both antibodies gave similar results. In younger animals (6-months), the level of APP immunoreactivity was relatively low (Fig. 3A, B). The cell soma of many of the larger neurons were marked by APP staining, but many neurons – both large and small – were unlabeled. By contrast, in the older animals (24-months), the intensity of APP immunoreactivity was markedly increased (Fig. 3C, D). The perikaryon of virtually every neuron was clearly labeled as were several smaller, presumably non-neuronal cells. We quantified the elevated levels of staining in the two different ages, calculating the staining intensity over the entire cell (nucleus plus pseudoplasm) with QuPath (Fig. 3E).

**Figure 3.**
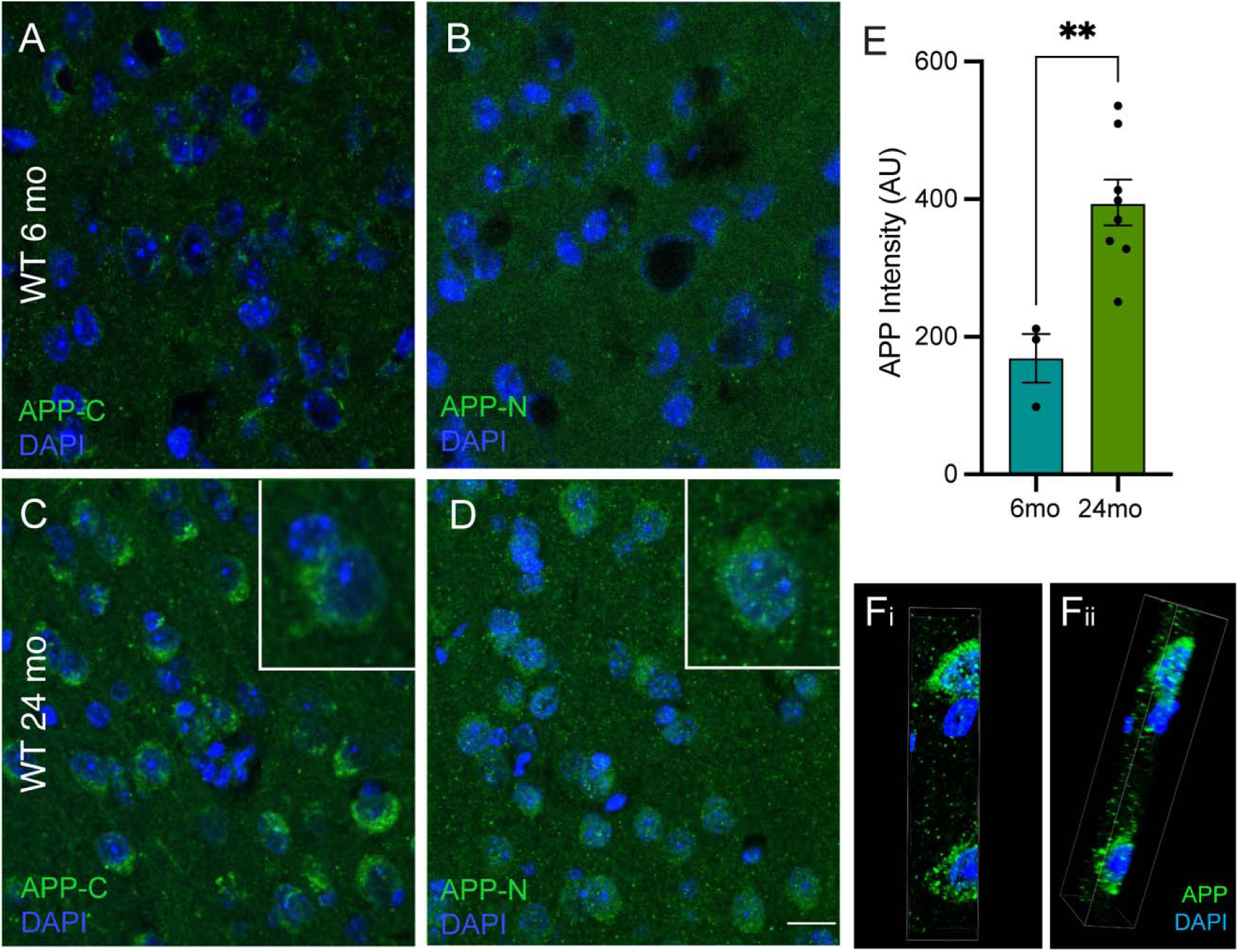
APP levels increase with age. (A) Neurons of layer V-VI from a 6-month-old wild type mouse prefrontal cortex immunostained for an APP C-terminal epitope (green) and counterstained for DAPI (blue). (B) An adjacent section from the same animal illustrated in (A) immunostained for an APP N-terminal epitope. Scale bar = 10 µm. (C) APP C-terminal immunostaining of a 24-month-old wild type mouse. (D) APP N-terminal immunostaining of a 24-month-old wildt ype mouse. In both (C) and (D) the inset is a cell from an adjacent region enlarged to show the details of the staining pattern. (E) APP intensity in young (teal) and old (green) neurons. Each point represents the average of all measured cells. Error bars represent standard deviation. **, p = 0.0045 by two tailed t-test (t = 3.758, df = 9). (F) Confocal image of 3 cells immunostained as in (C). (Fi) The Z-stack of confocal planes is viewed without rotation. (Fii) The same stack rotated to show the nuclear localization of the APP-C immunostaining.

Although the staining patterns of APP-C and APP-N were similar in most ways, there were also notable differences. These are most easily seen in the older brain sections where the APP staining intensity was stronger (Fig. 3C, D, and insets). The cytoplasmic APP-C signal was localized primarily in small puncta that could be found in both nucleus and cytoplasm. APP is not commonly considered to be a nuclear protein. We therefore validated its presence in the nucleus by confocal imaging (Fig. 3Fi and Fii). Although the APP-N antibody still clearly labeled the puncta seen with the APP-C antibody, we found the APP-N signal to be more evenly dispersed throughout the soma. There was also a notable increase in the background labeling in the APP-N material. The reasons for these differences are uncertain. Both antibodies will recognize the full-length protein. However, only APP-C will recognize the APP intracellular domain (AICD), and only APP-N will recognize the secreted sAPP fragments released following α-, β-, or η-secretase cleavage. As the two antibodies gave nearly identical patterns of labeling, we chose to use the knockout validated APP-N antibody for our quantitative work.

### Correlation of APP levels with DNA damage

The data thus far show that both DNA damage and intracellular APP levels increase with age. To determine whether APP accumulation was a response to DNA damage, thus implying a causal relationship between the two, we examined the average level of APP immunostaining intensity as a function of the level of DNA damage detected by the TUNEL reaction (Fig. 4A). Each point in this graph represents the average APP and TUNEL signals for all the measured cells in a single animal. Focusing solely on the data from the older animals, we performed a linear regression analysis. The line drifts upward, but the slope is not significantly different from zero (Dfn=1, DFd=6, p=0.4856).

**Figure 4.**
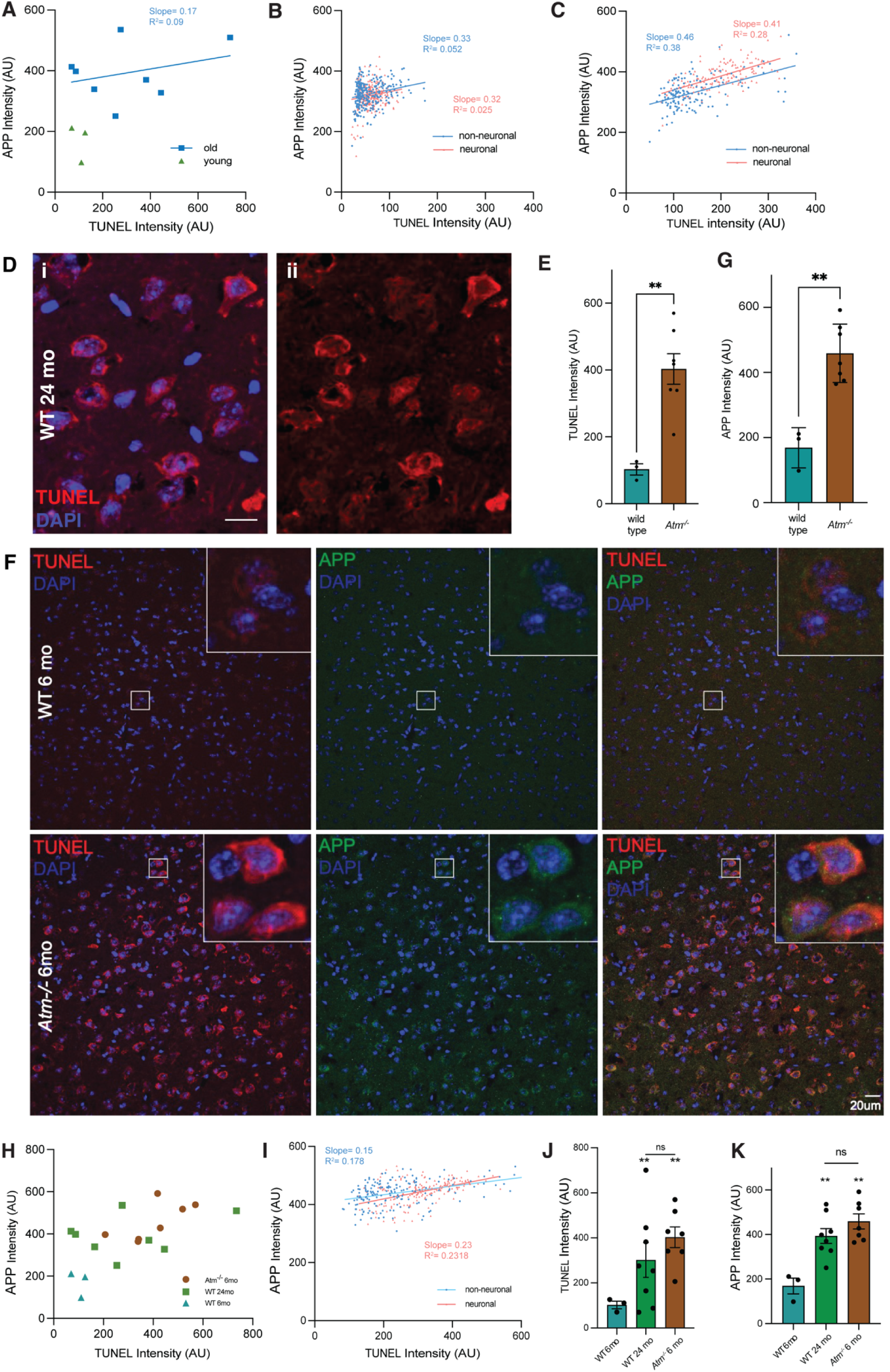
APP and TUNEL staining intensity are correlated. (A) APP-N immunostaining intensity plotted as a function of TUNEL staining intensity for layer V-VI neurons in frontal cortex of 6- (green) and 24-month (blue) wild type mouse. Data points represent an average value for all cells measured in one half brain. Slope = 0.17. The r^2^ value was 0.09. (B) Analysis of individual neuronal (pink) and non-neuronal (blue) cells from a single section of a 6-month-old wild type mouse (r^2^ values for neuronal and non-neuronal cells were 0.02527 and 0.05205, respectively). (C) Analysis of individual neuronal (pink) and non-neuronal (blue) cells from a single section of a 24-month-old wild type mouse. The r^2^ values for neuronal and non-neuronal cells were 0.28 and 0.38, respectively. (D) Neurons in a 6-month-old *Atm^−/−^*mouse stained with TUNEL and counterstained with DAPI. Scale bar = 10 µm. (E) Average TUNEL staining intensity for neurons of both wild type and *Atm^−/−^* genotypes. Data points represent an average value for all cells measured in one half brain. Error bars represent standard error. **, p = 0.0045 by two tailed t-test (t = 3.758, df = 9). (F) Cortex from a 6-month-old and a 24-month-old adult *Atm−/−* mouse labeled with TUNEL (red), immunostained for APP-N (green), and counterstained with DAPI (blue). (G) Intensity of APP immunostaining is significantly stronger in neurons of 6-month-old adult *Atm^−/−^* than in wild type mice. Error bars represent standard error. **, p = 0.001 by two tailed t-test (t = 5.034, df = 8). (H) Summary plot of APP as a function TUNEL staining in 6- and 24-month-old wild type, and 6-month-old *Atm^−/−^* mice. The slope of the old mice (green squares, replotted from (A) is 0.13 (r^2^ = 0.09). The slope of the *Atm^−/−^*mouse is 0.51 (r^2^ = 0.47). (I) Analysis of individual neuronal (pink) and non-neuronal (blue) cells from a single section of a 6-month-old *Atm^−/−^*mouse (r^2^ values for neuronal and non-neuronal cells were 0.2318 and 0.1780, respectively). (J) Average DNA damage (TUNEL staining) in the three genotypes (**, p ≤ 0.01 compared to 6-month wild type; 24-month wild type and 6-month *Atm^−/−^* were not significantly different (p = 0.2983). Error bars represent standard error. (K) Average APP immunostaining intensity in the three genotypes (**, p ≤ 0.01 compared to 6-month wild type); 24-month wild type and 6-month *Atm^−/−^* were not significantly different (p = 0.1910). One-way ANOVA with multiple comparison. Error bars represent standard error.

However, using the average of all cells in this way loses information. When we repeated the analysis using individual cells, the significance of the relationship was revealed. For these analyses we worked with data from single images; there were minor variations from one image to the next, but the pattern was the same throughout. We found that for both neurons and non-neuronal cells (large and small nuclei), APP levels increased linearly with the extent of DNA damage measured by the TUNEL reaction. The finding was consistent in both young (Fig. 4B) and old (Fig. 4C) samples. In the younger animals, the slopes of the lines for both neurons and non-neuronal cells had a positive slope, but the r^2^ values were low (0.06 or less) since the points were clustered at low TUNEL scores. In the older animals, the relationship was more robust. The slopes of neuronal and non-neuronal cells were greater (0.41 and 0.46, respectively) as were the r^2^ values (0.28 and 0.38 respectively). Note that in young animals, the Y-intercepts of the neuronal and non-neuronal cells were nearly identical, while in the older animals, even though the slopes were nearly equal, the Y-intercept of the neuronal line was higher than that for the non-neuronal cells (Fig. 4C).

The strong nature of the correlation between the levels of APP and DNA damage is compelling but does not prove a causal relationship. To test whether DNA damage was the driver of this phenomenon, we turned to mice deficient in the ATM (ataxia-telangiectasia mutated) kinase. ATM is a key component of the DNA damage repair pathway and in its absence, the normal process of DNA repair is compromised, both in vivo and in vitro (Shiloh, 2014). Inducing DNA damage in this independent way allowed us to query the APP response of cortical neurons in the absence of any non-specific effects of aging. We processed thirteen half brains from eight different 6-month-old animals homozygous for the *Atm^tm1Bal^*allele of the *Atm* gene. This allele carries a mutation that interrupts the kinase domain, leading to a catalytically inactive protein (Xu et al., 1996).

TUNEL labeling revealed a picture very similar to that found in 24-month wild type animals (Fig. 4D), with signal found in both nucleus and cytoplasm. Compared to age-matched wild type, the level of DNA damage was significantly higher in the mutant brains (Fig. 4E). Next, we asked whether there was any difference in the levels of APP in *Atm^−/−^*neurons compared to wild type. Both qualitatively (Fig. 4F) and quantitatively (Fig. 4G) the levels of APP were increased significantly in the absence of ATM activity. One curious feature of the APP staining was that in the 24-month wild type animals, puncta of immunoreactivity appeared in the nucleus (Fig. 3D, inset) whereas in the *Atm^−/−^*neurons, the increased APP signal was almost exclusively cytoplasmic (Fig. 3F).

The quantitative relationship between TUNEL and APP signals was intriguing. Plotting the data from all animals as in Figure 4A illustrated how the *Atm^−/−^* points tended to cluster in the same region of the graph as did the older wild type animals (Fig. 4H). Unlike the rather shallow slope to the curve of the old wild type animals, however, the average APP/TUNEL slope was steeper (0.51 compared to 0.13) and the relationship was more robust (r^2^ = 0.47 for the 6-month *Atm^−/−^* neurons; 0.09 for the old wild type). In any one animal (Fig. 4I), the relationship (slope and relative position of the neuronal and non-neuronal cells) resembled the curves for the old wild type animals except that the neuronal line was not displaced upward from that of the non-neuronal cells. Comparison of the average levels of DNA damage (Fig. 4J) and APP immunostaining (Fig. 4K) validates that there is significantly more DNA damage and more APP immunoreactivity in both young ATM-deficient and old wild type animals relative to the levels found in young wild type.

### Mouse APP increase does not lead to plaque formation but axon damage

Elevated levels of APP protein, as seen in *Atm^−/−^* and aged wild type mice, mimics the pathology seen in Alzheimer’s disease and could potentially lead to the formation of amyloid deposits. Secretase cleavages (β and γ) of mouse APP produce a peptide that is homologous to human Aβ, but the mouse peptide is not known to form extracellular amyloid deposits. For this reason, transgenic mouse models of human AD that do form plaques carry human APP transgenes in addition to, or in replacement of, the endogenous mouse APP gene (Sturchler-Pierrat et al., 1997; Richardson and Burns, 2002; Manabe and Saito, 2025). Nonetheless, to test the relationship of the elevated APP to the pathophysiology of AD, we performed immunohistochemistry of 6-month, 24-month, and *Atm^−/−^* mice sections with the 6E10 monoclonal antibody that recognizes both mouse and human Aβ (Fig. 5). No extracellular deposits of Aβ plaques were detected in any of our samples. When co-labeled with NeuN to visualize neurons, we did observe increased intracellular 6E10 staining in both 24-month-old wild type mice and 6-month-old *Atm^−/−^* neurons (Fig. 5A, insets). The intracellular 6E10 staining was stronger in the 24-month sample than in the *Atm^−/−^* mice. We then verified this finding using western blot analysis. We found that the 25 kDa 6E10 band (a species of Aβ oligomer) was more abundant in the 24-month wild type mice (Fig. 5B and C). Lower molecular weight bands corresponding to the Aβ monomer (~4 kDa) and smaller oligomers (~8-20 kDa) were absent from our samples.

**Figure 5.**
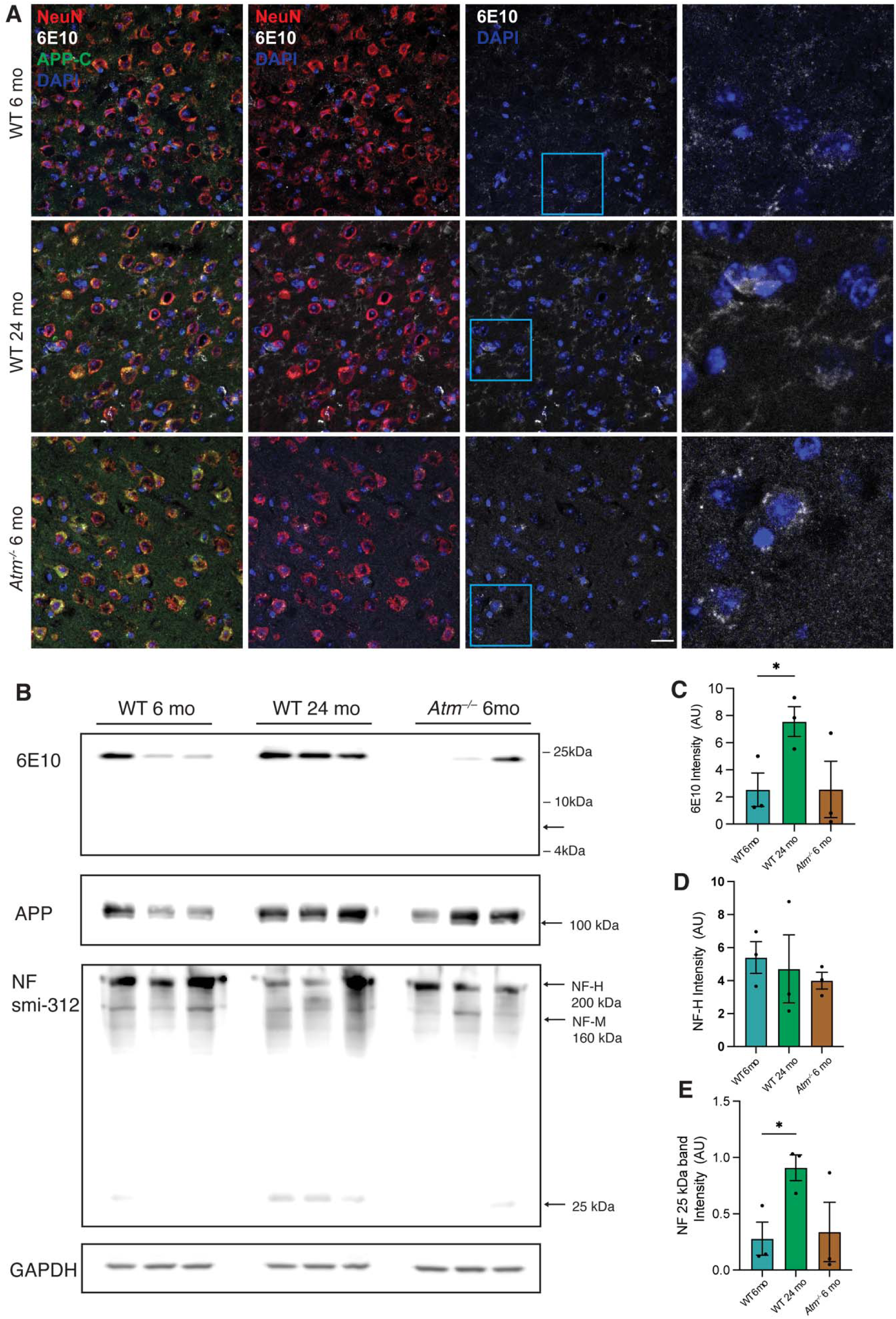
DNA damage and APP do not induce β-amyloid deposits in mouse but do increase evidence of axon damage. (A) Representative images of the cortex of a 6-month wild type mouse, a 24-month wild type and a 6-month *Atm^−/−^* mouse labeled with NeuN (red), 6E10 (gray), APP C-terminal (green) antibodies and counterstained with DAPI (blue). Insets showed the region in the blue box. Scale bar = 20 µm. (B) Western blot of 6E10, N-terminal APP, and Neurofilament smi-312 of lysates from young wild type, old wild type and *Atm^−/−^* brains. (C) Quantification of the 25 kDa bands of 6E10 shown in (B). (D) Quantification of the NF-H 200 kDa bands shown in (B). (E) Quantification of the neurofilament 25 kDa bands shown in (B). Data from (C), (D) and (E) were all normalized to their respective loading controls. All error bars represent standard error. *, p < 0.05 (One-way ANOVA with multiple comparison).

We also tested a second marker of AD pathogenesis, the presence of degraded neurofilament protein. Although the difference did not reach statistical significance in our samples, we found a notable decrease of neurofilament heavy and medium chain species in the 24-month-old wild type mice and *Atm^−/−^*mice in addition to an increase in the 25 kDa neurofilament breakdown product (Fig. 5B, D and E). These changes presumably reflect early events of axonal loss or degeneration (Schlaepfer et al., 1985; Gou and Leterrier, 1995; Yuan et al., 2017; Cao et al., 2020; Yuan and Nixon, 2021).

## DISCUSSION

Our work describes the inter-relationships between three forces that play a role in Alzheimer’s disease: age, DNA damage, and the amyloid precursor protein (APP). Several important findings emerge from the study related to each of these. We show that DNA damage increases in neuronal cells as the organism ages. As early as six months (~20% of the life span of a mouse) low levels of broken DNA ends are already apparent, and by two years of age the levels increase nearly three-fold. This finding is similar to work done by other labs including our own (Katyal and McKinnon, 2008; Chow and Herrup, 2015; Maynard et al., 2015; Tse and Herrup, 2017; Gutta et al., 2024).

Utilization of the TUNEL technique to track DNA damage adds important information to the discussion. Most studies monitor DNA damage using immunostaining of DNA repair-related proteins such as γ-H2AX and 53BP1. All such assays are indirect in their nature because they detect the cellular response to DNA damage rather than the damage itself. In contrast, the TUNEL reaction enzymatically extends any free 3’ end of a DNA strand, and thus directly detects the presence of damage independent of the cell’s response to it. Not surprisingly, therefore, the staining patterns of Y-H2AX and TUNEL are different. Also as expected, the Y-H2AX immunostaining was more restricted to the nucleus than that of the TUNEL reaction. The fragments of cytoDNA, while rich in free 3’ ends and hence revealed by TUNEL are A-T-rich sequences that are short (~100 bp) and thus less likely to be decorated with histone proteins such as H2AX (Song et al., 2021). In microglial cells, DNA in the cytoplasm triggers a sterile immune response using cGAS/STING signaling. The consequence of cytoDNA in neurons is unknown. Its levels as detected by TUNEL are substantial, but nerve cells lack cGAS/STING. Whether there is a yet unrecognized cytoDNA response pathway or whether it accumulates because it cannot be efficiently cleared is unknown.

Given the interest in its Aβ breakdown product, it is surprising that searching the literature for papers that measured changes in the levels of full-length APP with age or Alzheimer’s disease uncovers very few studies. Single-cell databases track mRNA levels rather than protein, but these data rarely include age as an independent variable. In our study, correlated with the increase in DNA damage, we found a two-fold increase in the intracellular levels of APP as animals age from 6 months to two years. To detect the APP protein, we relied on two different antibodies, neither of which would detect the short Aβ peptide. The APP-C antibody targets an epitope in the cytoplasmic domain that is retained in the AICD. The epitope used to raise the APP-N antibody is listed as proprietary. By analogy with the 22C11 mouse monoclonal antibody, it would be expected to recognize amino acids in the region of the protein encoded by exons 2 and 3 and would thus recognize secreted forms of APP (created by α-, β-, or η-secretase cleavage) as well as the full-length protein.

Comparison of the 6- and 24-month wild-type staining patterns revealed that both nuclear and cytoplasmic APP increase. In the cytoplasm its localization was diffuse, but clear APP puncta were present that could represent vesicular or Golgi-localized APP. Unexpectedly, APP puncta were also present in the nucleus as confirmed by confocal-imaging. The APP intracellular domain (AICD) has been shown to translocate to the nucleus where it can affect gene expression (Gao and Pimplikar, 2001; Cao and Sudhof, 2004) and modulate DNA repair (Revol et al., 2023). The finding of nuclear APP-N puncta suggests that there may be additional undiscovered roles for full length or secreted APP fragments in the nucleus.

After secretase cleavage, endogenous mouse APP produces an Aβ-like peptide that does not spontaneously aggregate to form plaques. Knock-in or overexpression mouse models of AD rely on the expression of human APP to develop amyloid plaques in mouse brain (Sturchler-Pierrat et al., 1997; Richardson and Burns, 2002; Manabe and Saito, 2025). Consistent with this, the increased levels of mouse APP in mice with excessive DNA damage – caused either by the lack of DNA repair (*Atm^−/−^*) or the accumulation of damage with advanced age – are not sufficient to induce the formation of Aβ plaques. In addition, mouse Aβ does not tend to aggregate compared to human Aβ(Xu et al., 2015). Intracellular 6E10 immunoreactivity does accumulate in neurons, suggesting that APP processing is following the path found in human AD, a suggestion that is strengthened by the observed increased breakdown of the neurofilament protein with age or ATM deficiency. We note with interest that despite the increase in both APP and DNA damage, ATM-deficient mice showed less evidence of axonal damage than did old wild type mice. This suggests that age increases neuronal vulnerability in dimensions other than DNA damage or levels of APP protein. It is intriguing to imagine that these other dimensions may help explain why diverse diseases such as ataxia-telangiectasia and Alzheimer’s disease differ in clinical outcomes despite sharing similarities in their pathogenesis.

Both APP and DNA damage increase with age, but our observations in the *Atm^−/−^* mice are the first to offer evidence that DNA damage itself drives the increase in APP. The levels of APP in 6-month-old *Atm^−/−^*neurons were as high as those found in 24-month-old wild type, providing strong evidence in support of the causal direction of the relationship. As the *Atm^−/−^*mice were age-matched to the 6-month wild type animals, the data do not support an alternative model where age itself, or an independent age-related factor, is necessary for the APP increase. The steeper slope of the curve shown in Figure 4H suggests that when DNA damage is precociously increased, the APP response is more robust. This too supports a model in which APP levels are responsive to the levels of DNA damage, just as they are to other cellular stressors (Ma et al., 2023). An important caveat to this logic is that DNA damage may drive the entire aging process, not just APP levels. The occurrence of symptoms of progeria found with mutations in DNA repair proteins supports this broader view. Altered expression of APP may thus be one of many symptoms of an aging brain, albeit an important one for the pathogenesis of Alzheimer’s disease.

## AUTHOR CONTRIBUTIONS

Conceptualization- F.M. and K.H.

Methodology -E.B., F.M., and K.H.

Investigation-E.B., F.S., M.B., S.W., N.D, E.Z., R.K., and F.M.

Visualization-E.B. M.B., S.W., N.D., E.Z., and R.K.

Software-E.B., G.G., J.M., and F.M.

Supervision-F.M. and K.H.

Writing-E.B., F.M., and K.H.

Funding Acquisition-K.H.

All authors have read and agreed to the published version of the manuscript.

## ACKNOWLEDGMENTS

Funding: University of Pittsburgh, the NIH (NS120922 and AG069912), Pennsylvania Department of Health (Project Number:4100087331), and the University of Pittsburgh David C. Frederick Honors College Research Fellowship.

## Institutional Review Board Statement

Not Applicable.

## Informed Consent Statement

Not applicable.

## Data Availability Statement

All data will be made available to qualified researchers upon request to the corresponding author.

